# Transcriptomic Profiling Reveals Claudin 18.2 as a Diagnostic Biomarker of Ménétrier’s Disease and the Role of Hedgehog Signaling in Pathogenesis

**DOI:** 10.1101/2023.11.03.565570

**Authors:** Miyoung Shin, Tryston Gabriel, Robert J. Coffey, Won Jae Huh

## Abstract

Both Ménétrier’s disease (MD) and juvenile polyposis syndrome (JPS) are rare premalignant conditions that can lead to gastric cancer development. MD is an acquired disease without known causative mutations. MD patients are characterized by an increased expression of EGF receptor (EGFR) ligand and transforming growth factor alpha (TGF-α) in the stomach. JPS is inherited in an autosomal dominant pattern and is caused by *BMPR1A* or *SMAD4* mutations. It is characterized by multiple polyps throughout the gastrointestinal tract along with certain *SMAD4* mutations that can result in gastric polyposis. Although there are many distinct clinico- endoscopic and histopathologic features that differ between the two diseases, they also share similar features that often lead to misdiagnosis. This study aimed to identify markers that can help distinguish MD from JPS and to better understand the pathogenesis of MD by comparing differential gene expression patterns.

Upon examination of MD and JPS microscopically, we found almost all cases have patchy areas mimicking each other, making it difficult to make a correct diagnosis with histopathologic examination alone. Comparative analysis between MD and JPS using ingenuity pathway analysis (IPA) revealed both common and differential gene signatures. Common gene signatures included estrogen receptor signaling, integrin signaling, mTOR signaling, and others, which may be responsible for histopathologic similarities. Among differential gene signatures, we found that *claudin 18* (*CLDN18*) is upregulated in MD and confirmed that CLDN18.2 (isoform of CLDN18) protein expression is higher in MD than JPS by immunohistochemistry. Comparative analysis between MD and normal control revealed the hedgehog (Hh) signaling pathway is upregulated in MD. Treatment with a hedgehog pathway inhibitor partially rescued the histopathologic phenotypes in a MD mouse model.

The current study provides valuable insight into the potential underlying mechanism of why MD and JPS show similar clinico-pathologic features. We also identified a diagnostic marker CLDN18.2 that can help distinguish MD from JPS, genetically. Furthermore, it also shows that Hh signaling plays an important role in the pathogenesis of MD and can function as a potential therapeutic target.

## INTRODUCTION

Ménétrier’s disease (MD), also known as protein losing hypertrophic gastropathy, is a rare premalignant gastric disorder. It is thought to be acquired and there are no known genetic mutations associated with the disease. MD is characterized by giant gastric rugal folds in the body and fundus, decreased acid secretion, increased gastric mucus production, and hypoalbuminemia secondary to protein loss that leads to systemic edema. Microscopically, it shows massive foveolar hyperplasia which causes increased secretion of mucin and a decrease in the number of parietal cells which secrete acid in the stomach (Coffey, Washington et al. 2007, Huh, Coffey et al. 2016). It has been shown that EGF receptor (EGFR) ligand transforming growth factor alpha (TGF-α) expression is increased in the stomachs of MD patients. Transgenic mice that overexpress TGF-α have been shown to present with similar histopathologic features of MD (Dempsey, Goldenring et al. 1992). A clinical trial using EGFR inhibiting antibody cetuximab showed it is the first effective medical therapy for MD (Fiske, Tanksley et al. 2009). During the initial patient selection, almost half of the original 48 patients that had been previously diagnosed with MD were found to have been misdiagnosed. The most common features mistaken were polyposis syndromes with gastric involvement or gastric hyperplastic polyps. Among the polyposis category of disease, juvenile polyposis syndrome (JPS) was the most common disease misdiagnosed as MD (Rich, Toro et al. 2010). JPS is a rare premalignant condition which causes numerous juvenile polyps in the gastrointestinal tract. It is caused by mutations in *SMAD4* or *BMPR1A* genes, and it follows an autosomal dominant inheritance pattern (Dal Buono, Gaiani et al. 2022). The most common area of polyposis is the colorectum; however, mutations in the *SMAD4* gene have been associated with massive gastric polyposis, which could mimic MD endoscopically and microscopically (Burmester, Bell et al. 2016, Bernshteyn, Bhutta et al. 2022, Jelsig, Qvist et al. 2022).

In this study, we aimed to identify diagnostic markers to help distinguish MD from JPS along with MD-specific signal pathways that enable a better understanding of the pathogenesis and can function as potential therapeutic targets. We report that claudin18.2 is highly expressed in MD when compared to JPS and can be used as a diagnostic marker. We also found that Hh signaling is upregulated in MD. By treating the mouse model with a Hh inhibitor, we saw a partial rescue of the phenotype, suggesting this pathway has an essential role in the pathogenesis of MD.

## MATERIALS AND METHODS

### Human samples and RNA isolation

Frozen stomach tissues from MD and JPS patients were obtained from the prior clinical trial of cetuximab treatment for MD patients (Fiske, Tanksley et al. 2009). Stomach tissues from normal individuals were obtained from the Cooperative Human Tissue Network (CHTN) - Western Division at Vanderbilt University Medical Center. Total RNA was extracted with RNeasy plus Mini Kit (Qiagen, US) from frozen tissue according to the manufacturer’s protocol.

### RNA-sequencing and gene expression analysis

The sequencing was performed on the Illumina platform (PE150, Q30 more than >80%), and the fastq files were aligned with BWA to produce bam files. The aligned dataset was uploaded on the Partek^®^ Flow^®^ software (Partek LLC, US) server for further analysis. A QC/QA post-alignment module was used to exclude low quality samples. Principal component analysis (PCA) and hierarchical clustering were used for classification of the samples. To identify differentially expressed genes, we performed the DESeq2 algorithm under the Partek^®^ Flow^®^ pipeline with cutoff value of fold change (FC) more than >2 or less tan <-2. Core analysis and analysis by means in the Ingenuity ® Pathway analysis (IPA, Qiagen) software were utilized to identify the unique MD canonical pathways, upstream regulators, and biological processes different from NL or JPS groups.

### Quantitative PCR

Total RNA was extracted from the frozen human stomach tissues using the RNeasy plus Mini Kit (Qiagen, US) according to the manufacturer’s protocol. Two μg of total RNA from each sample were used for cDNA synthesis using the iScript Advanced cDNA Synthesis kit (Bio-Rad, CA).

SsoAdvanced Universal SYBR^®^ Green Supermix (Bio-Rad, CA) was used for visualizing the amplicons of target genes: *CDX2* (Forward: 5’-CAACCTGGCGCCGCAGAAC-3’; Reverse: 5’- GAGTGGGGCGCCATACGCTGC-3’), *HES1* (Forward: 5’ -CCTGTCATCCCCGTCTACAC-3’; Reverse: 5’-CACATGGAGTCCGCCGTAA-3’), *GLI1* (Forward: 5’- AACGCTATACAGATCCTAGCTCG-3’; Reverse: 5’- GTGCCGTTTGGTCACATGG-3’), *RPS18* (Forward: 5’- CGCTGAGCCAGTCAGTGTAG-3’ Reverse: 5’-CGCTTCGGCAGCACATATACTA-3’). The quantification of amplicon was detected by CFX Opus 96 Real-Time PCR system (Bio-Rad, CA). The relative quantification of amplicon was normalized with ribosomal protein 18 (RPS18) and used to calculate the relative expression of target genes following the Livak-Schmidt (2^-ΔΔCt^) method.

### Mice

All animal husbandry and experiments were approved by the Institutional Animal Care and Use Committees at Yale University (2021-20422) and in accordance with U.S. Government Principles for the Utilization and Care of Animals Used in Research, Teaching and Testing. MT- TGF-α mice (2 - 4 months old) were treated with sonidegib (LDE225; NVP-LDE225) (5 mg/kg; Med Chem Express, HY-16582A) in 10% ethanol/90% corn oil (Sigma Aldrich, C8267) or 10% ethanol/90% corn oil intraperitoneally every other day. Mice were sacrificed on day 21 and stomachs were dissected and fixed in 10% formalin or frozen fresh in liquid nitrogen.

### Immunoblot analysis and Immunofluorescence (IF) microscopy

Protein was extracted by 1x RIPA lysis buffer (10x, stock solution, EMD Millipore, MA) from mouse stomach tissue, separated by 4-12% gradient Bis-Tris SDS-PAGE gel (Invitrogen, CA), and transferred to a polyvinylidene difluoride membrane (PVDF) using an iBlot2 gel transfer device (Invitrogen, CA). The membrane was blocked with 3% Bovine Serum Albumin (BSA) in 1x TBST (TBS with 0.1% Tween-20) solution for an hour and incubated with the following primary antibodies: GLI1 (1:1000, Bioss, Bs-1206R), beta-Actin (1:3000, Proteintech, 20536-1- AP) at 4 °C for overnight, and then incubated with horseradish peroxide (HRP)-conjugated secondary antibody (1:5000, Invitrogen, A16035) for an hour at room temperature. Chemiluminescence HRP substrate (Thermo Scientific, 34096) was used for detection by ChemiDoc XRS+ imaging system (BIO-RAD, CA). Formalin-fixed paraffin-embedded (FFPE) sections were used for immunofluorescence staining with ulex europaeus agglutinin I (UEA I), Fluorescein (1:500, Vector Laboratories, FL-1061) and griffonia simplicifolia-II (GS-II), Alexa Fluor 594 conjugate (1:500, Invitrogen, L21416). Fluorescent signals were visualized using a fluorescence microscope (KEYENCE, BZ-X800, Japan).

### Histology, immunohistochemistry (IHC), and Scoring

FFPE blocks were obtained from the prior clinical trial of cetuximab treatment in MD patients (Fiske, Tanksley et al. 2009). Hematoxylin and eosin (H&E) stain and immunohistochemical stain for CLDN18.2 (LS-B16145, LS Bio, MA) were performed by Yale Pathology Tissue Services (YPTS). Whole slide images were acquired by Motic EasyScan system (Motic, TX). IHC and IF images were analyzed with QuPath software (ver 0.4.3) (Bankhead, Loughrey et al. 2017). H-score was calculated by adding 3x % of strongly stained positive cells, 2x % of moderately stained positive cells, and 1x % of weakly stained positive cells, giving results in range 0 to 300.

### Statistics

GraphPad Prism version 10 (GraphPad Software, CA) was used for all statistical analyses. All data is presented as means ± SEM. Grouped two-way analysis of variance (ANOVA) or unpaired student t-test was used for the statistical analyses between two groups or among the three groups.

## RESULTS

### Histopathologic features of MD and JPS

In the cetuximab clinical trial of MD, gastric polyposis was the most common disease entity that was misdiagnosed as MD. Among gastric polyposis, JPS was most commonly misdiagnosed as MD. Despite MD and JPS both exhibiting foveolar hyperplasia, it has been generally perceived that each disease has its own distinct histopathologic features. MD shows massive foveolar hyperplasia with corkscrew morphology, decreased number of parietal cells, and vertical growth of smooth muscle in the lamina propria, but glandular architecture is largely maintained (Figure 1, upper left). In contrast, JPS shows foveolar hyperplasia, cystically dilated glands and expanded lamina propria with edema, but number of parietal cells is not decreased (Figure 1, lower left). When we closely examined histopathologic samples of MD (3/7) and JPS (2/4), we observed areas that display nearly identical morphology (Figure 1, right), which may lead to misdiagnoses.

**Figure 1.**
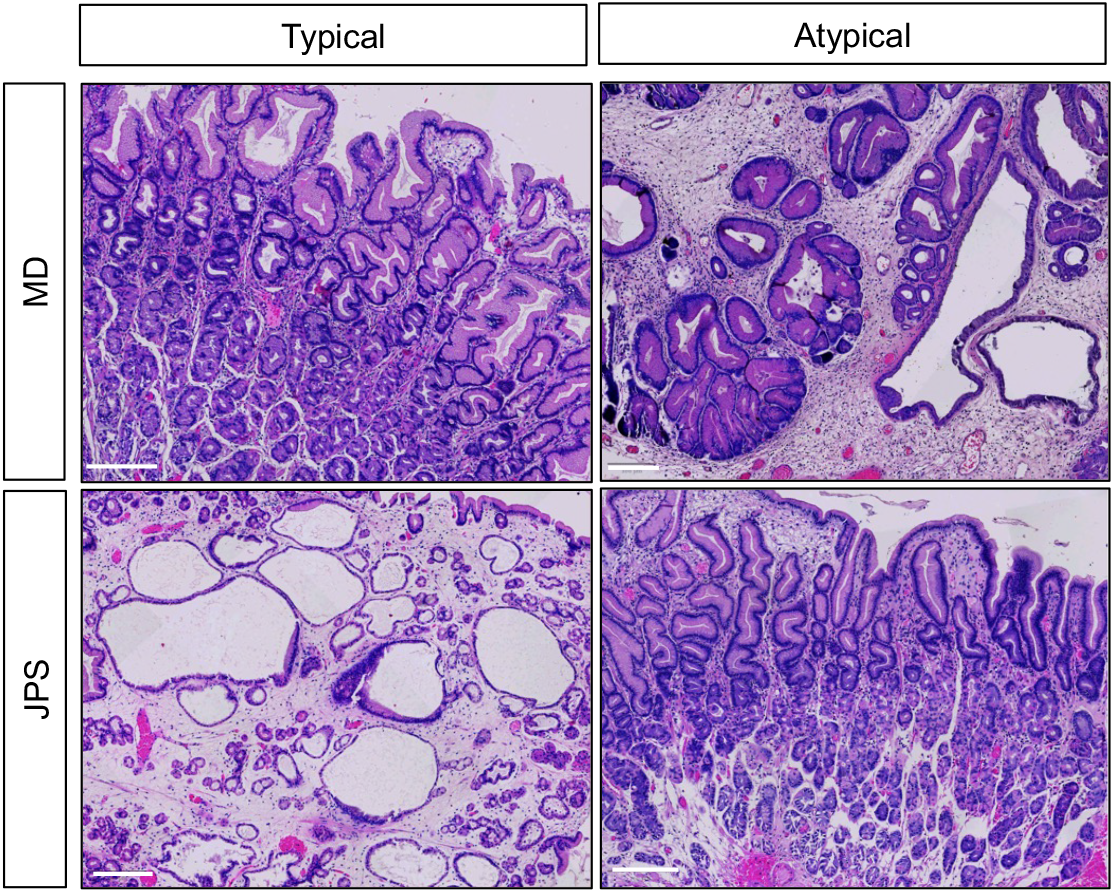
Histopathological features of MD and JPS. Hematoxylin and eosin (H&E) stains of MD (top) and JPS (bottom) show both areas with typical histopathologic features (left) and areas with atypical histologic features mimicking each other (right). MD usually presents massive foveolar hyperplasia with corkscrew morphology, decreased number of parietal cells, and vertical smooth muscle growth in the lamina propria. However, overall glandular architecture is maintained. JPS shows cystically dilated glands with foveolar hyperplasia, and expanded lamina propria with edema. Most MD and JPS cases showed areas of histopathologically mimicking each other, which can lead to misdiagnosis. The scale bar represents 200 µm.

### Mining the RNA-sequencing datasets in MD, JPS and Normal stomach tissues

To find diagnostic markers that can distinguish MD from JPS and to better understand the pathogenesis of MD, we performed RNA sequencing (RNA-seq) using gastric mucosa from three MD patients, three JPS patients, and three normal individuals (NL). Quality analysis revealed that two samples (JPS2 and NL2) have low average coverage (less than 10x average coverage) and low read counts (Figure 2A and 2B). These two samples were also outliers in the similarity and correlation analysis (Figure 2C). These findings suggest that JPS2 and NL2 samples are suboptimal in quality and data from these samples will not be reliable. Therefore, we decided to exclude them from further analysis. Despite excluding these two outliers, the principal component analysis (PCA) revealed a lack of similarities across the three phenotypes, with two JPS samples not segregating from each other (Figure 3A). Considering the characteristics of MD including the increased expression of TGF-alpha (TGF-α) (Coffey, Washington et al. 2007), we added *TGFA* as an additional attribute for further analysis that led to a more uniform and defined segregation among different phenotypes (Figure 3B). It also showed each group categorized together in the hierarchical clustering when each sample was grouped by TGFA read counts: MD samples in high TGFA (>500 of average read counts), JPS samples in mid TGFA (200-500 of average read counts), and NL in low TGFA (<200 of avg. read counts) as shown in Figure 3C.

**Figure 2.**
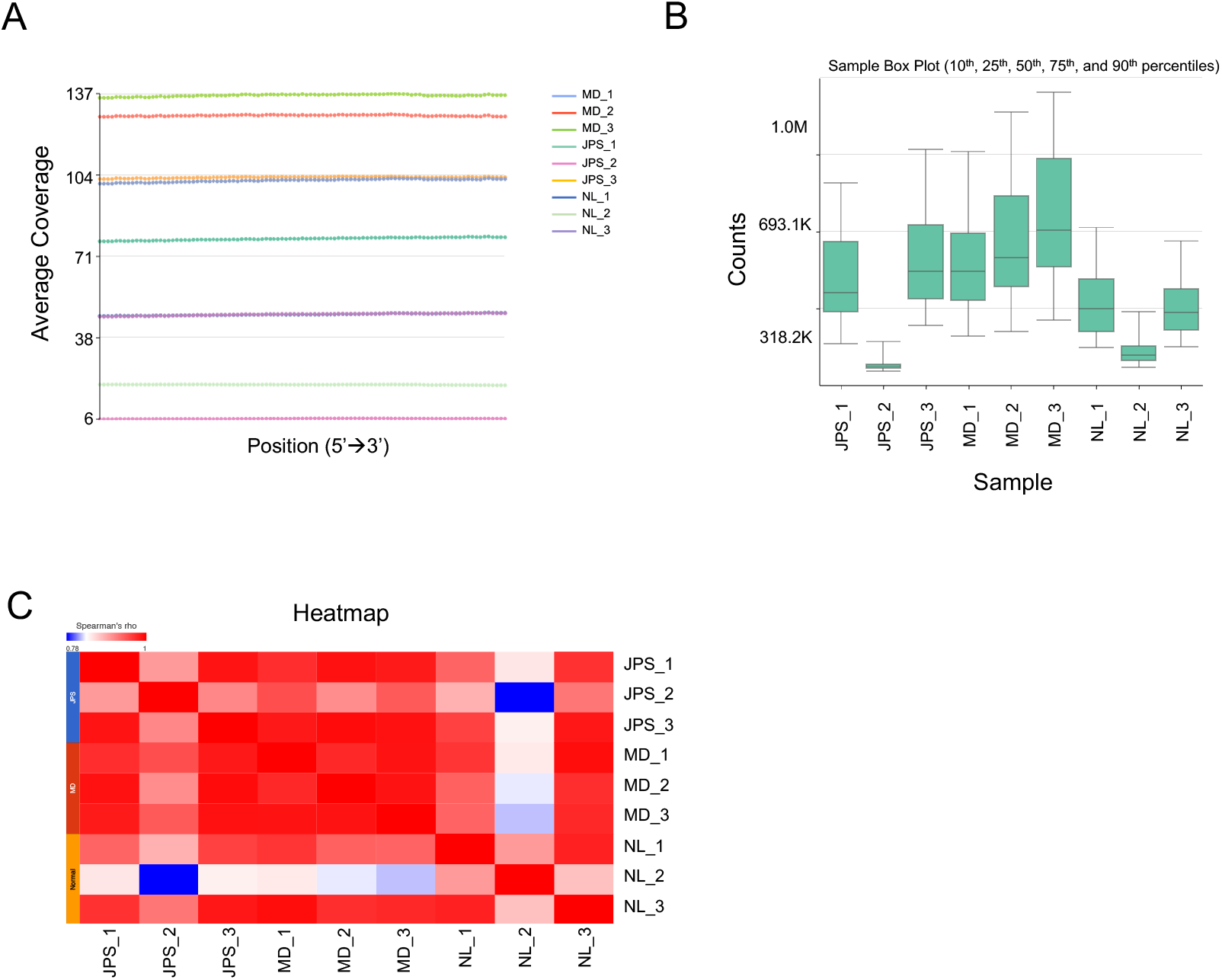
Quality analysis of RNA-sequencing (RNA-seq) dataset. (A) Summary of coverage from nine transcriptomic datasets presents the average coverage of each sample. JPS 2 and NL 2 show the lowest coverage. (B) The read counts of nine samples reveal the lowest values in JPS 2 and NL 2. (C) Heatmap from the Spearman’s correlation analysis among nine samples highlights that JPS 2 and NL 2 are outliers. Detailed information of the heatmap can be found in supplementary Table S1.

**Figure 3.**
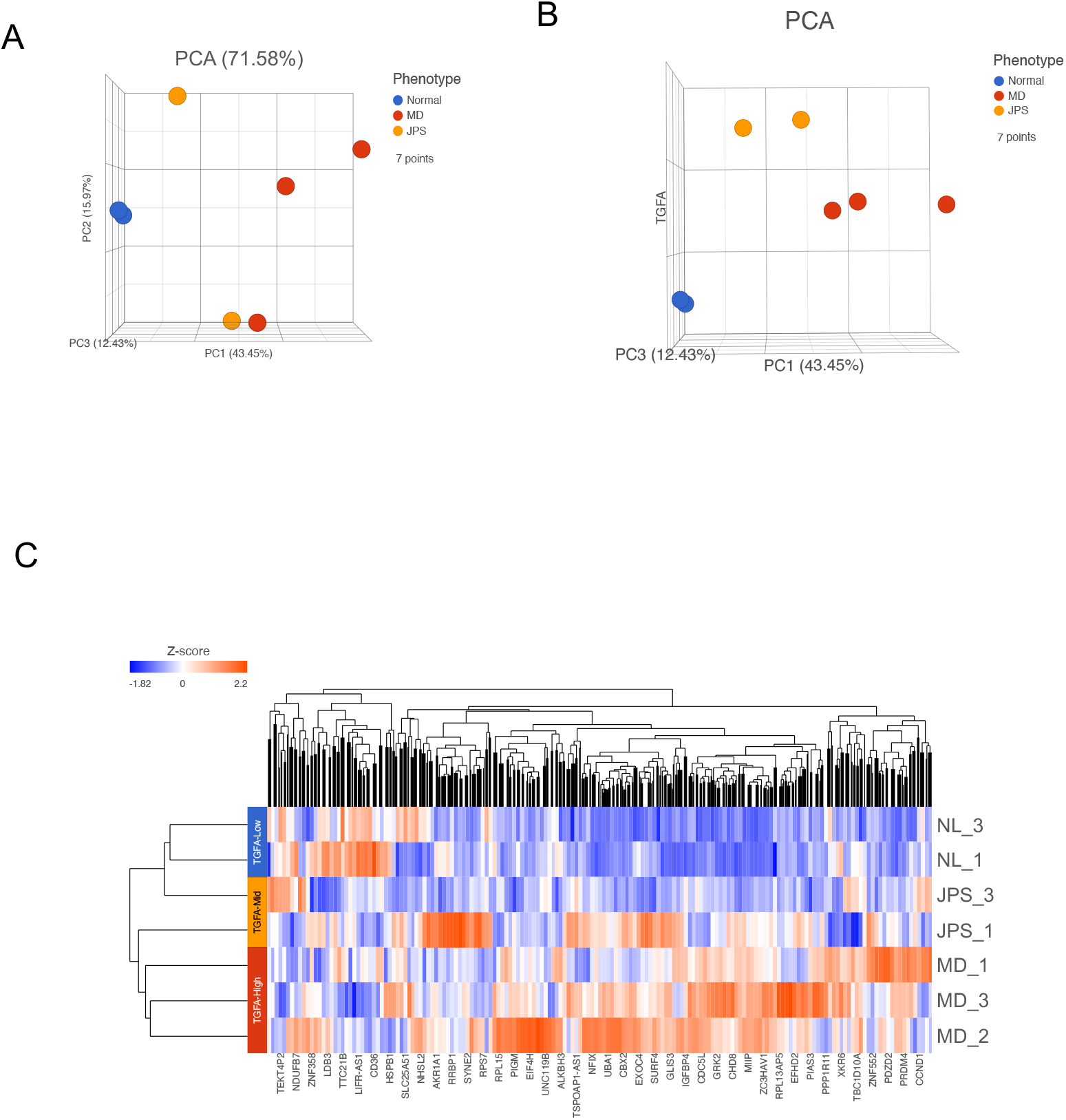
Clustering analysis of RNA-sequencing datasets. (A) Principal component analysis (PCA) of the nine samples across the three groups, Normal (blue), MD (red) and JPS (yellow) reveals that JPS samples are not segregated together. (B) Adding an additional attribute of transforming growth factor alpha (*TFGA*) expression made each phenotype segregated separately. (C) Heatmap with additional attribute of differential expression level of *TGFA* causes clustering among the same phenotype group (z-score range from – 1.82 to 2.2). TGFA-low: read counts <200, TGFA-mid: read counts from 200 to 500, and TGFA-high: read counts >500.

### Comparative analysis reveals commonly regulated genes and pathways between MD and JPS

We conducted a comparative analysis between MD and JPS using the DESeq2 algorithm under the Partek Flow pipeline with the genes expressed in MD or JPS exhibiting more than >1 of log2(Fold Change) compared to NL. We obtained 646 and 680 genes in MD and JPS, respectively. Among those genes, 487 genes were common between MD (75.3%) and JPS (71.6%). The pathway analysis performed using the common genes between MD and JPS revealed estrogen receptor signaling, integrin signaling, mTOR signaling and others were upregulated in the two disease groups (Figure 4A). Molecules in the common signaling pathway were involved in cell transformation, development of epithelial tissue, growth of embryo, and advanced malignant tumor, among others (Figure 4B).

**Figure 4.**
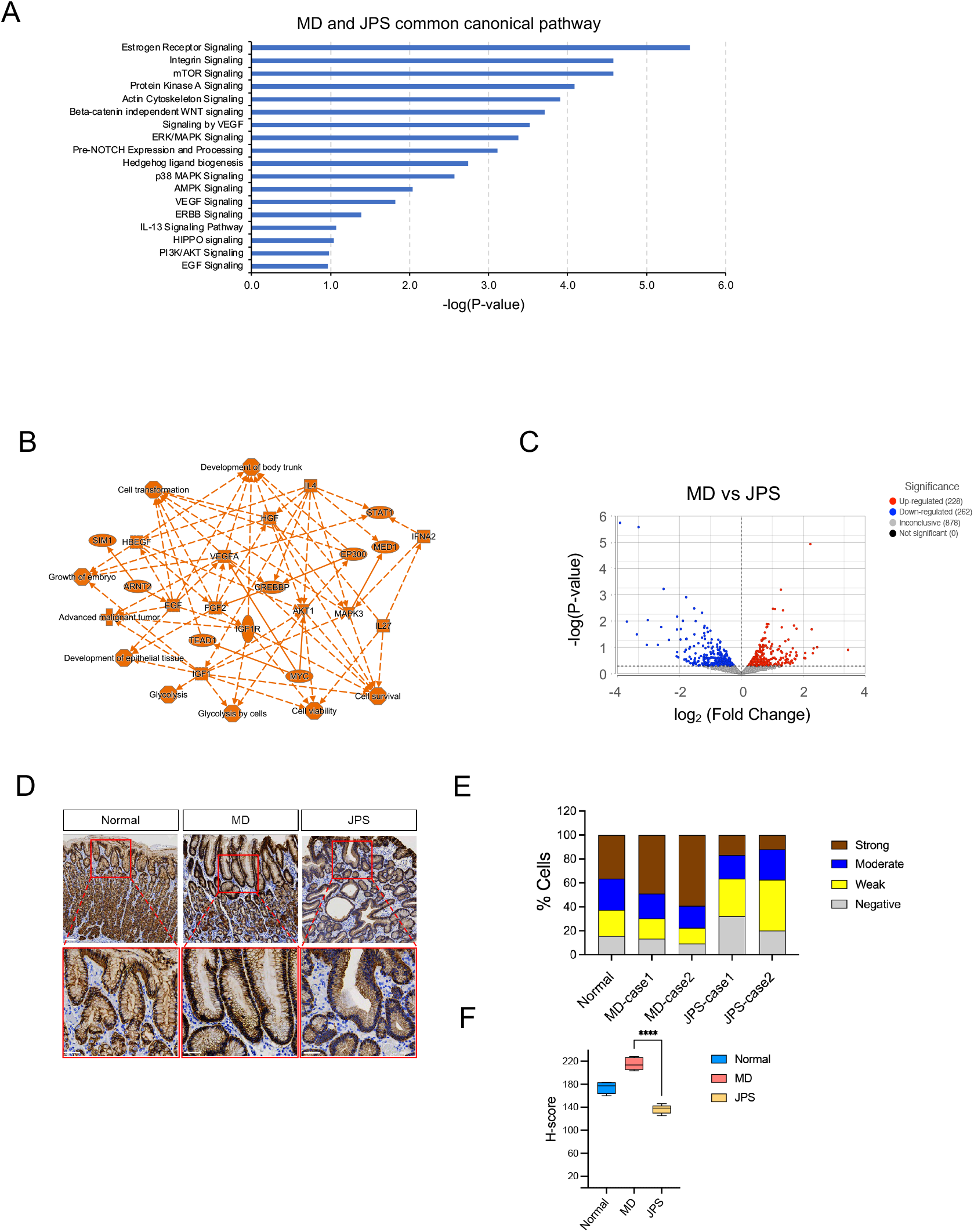
Comparative analysis between MD and JPS. (A) Ingenuity pathway analysis (IPA) with the genes that are expressed higher than normal in both MD and JPS groups (485 genes with log2(Fold Change) >1). Complete list of the genes can be found in the supplementary Table S2. (B) Graphical network summary of the common genes from MD and JPS. Orange-colored nodes and lines show predicted activation in the network. (C) Volcano plot of differentially expressed genes between MD and JPS (cutoff range: -1 < log2(Fold Change) < 1). (D) Immunohistochemistry (IHC) staining of Claudin 18 isoform 2 (CLDN18.2). Scale bars represent 200 μm. (E) Percentage of cells with different staining intensity of CLDN18.2 (negative, weak, moderate, and strong) in normal, MD, and JPS cases. (F) H-scores are significantly different between MD and JPS with grouped two-way ANOVA (**** means P value <0.0001). H-scores were calculated by the QuPath (ver. 0.4.3) software.

### Differential expression analysis between MD and JPS reveals that claudin18.2 is highly expressed in MD

Based on the differential gene expression, we aimed to identify MD biomarkers that can be used to distinguish MD from JPS presenting with similar histopathologic features. As visualized in the volcano plot of MD vs. JPS, we found 228 upregulated genes and 262 downregulated genes in MD (Figure 4C). We investigated the upregulated genes in MD utilizing IPA^®^ biomarker profiling and identified 68 therapeutic target genes (Supplementary Table S3). Among those target genes, we particularly considered claudin18 (CLDN18) as a potential diagnostic biomarker for MD since CLDN18.2 (CLDN 18 isoform 2) expression has been reported as a biomarker for targeted therapy of gastric cancer in several clinical studies (Dottermusch, Kruger et al. 2019, Lu, Wu et al. 2020, Kubota, Kawazoe et al. 2023). Immunohistochemical staining for CLDN18.2 was performed on stomach tissues from MD, JPS, and NL (Figure 4D) and images were analyzed with QuPath software. CLDN18.2 expression was highest in MD samples both in percent positivity and signal intensity (Figure 4E). This was quantified using the H-score, and MD group had significantly higher H-score than both JPS and normal groups (Figure 4F).

### Comparative analysis between MD and NL reveals signaling pathways enriched in MD

To better understand the pathogenesis of MD, comparative analysis between MD and Normal was performed and it resulted in 271 upregulated genes and 266 downregulated genes in MD (Figure 5A). We obtained extended candidate genes from the canonical pathway analysis and graphical summary module of IPA^®^ software, which showed the relationship between gene expression and its function in human disease (Figure 5B). We performed the canonical pathway analysis in the IPA software with the differentially expressed genes in MD compared to NL. From these differential genes, we found many signal pathways (blue bars) that were communal between MD and JPS as shown in Figure 4A. Of these, we focused on MD-specific pathways (orange bars): CDX, Notch, and Sonic Hedgehog (SHH), that were not found to be upregulated in JPS. We tried to validate several target genes out of the pathways enriched in MD using quantitative real time PCR (qRT-PCR): *CDX2*, *HES1*, and *GLI1* are the downstream effector molecules in the CDX gastrointestinal cancer signaling pathway, Notch signaling pathway, and SHH signaling pathway, respectively (Figure 5D). *CDX2* and *HES1* expression showed trends of upregulation in MD, but they were not statistically significant. Interestingly, the expression of *GLI1* and other hedgehog (Hh) signaling downstream target genes, such as *HHIP*, and *BOC* (data not shown) were significantly upregulated in MD when compared to JPS and NL.

**Figure 5.**
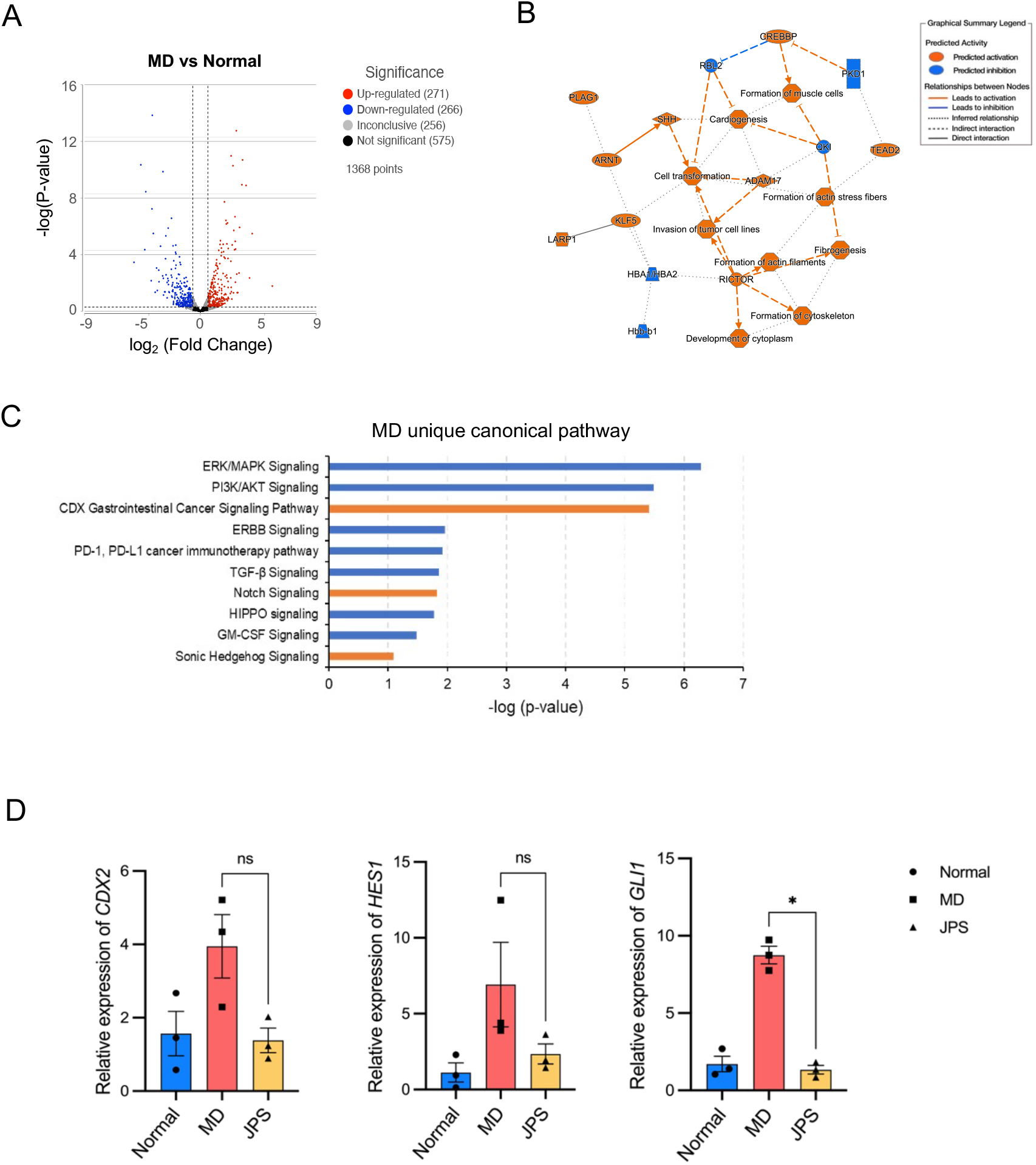
Differential expression analysis between MD and Normal. (A) Volcano plot of differentially expressed genes between MD and Normal groups (cutoff range: -1 < log2(Fold Change) < 1). Complete list of the genes can be found in the supplementary Table S4. (B) Graphical network summary of MD-unique gene expression. Orange-colored nodes and lines show predicted activation, and blue-colored nodes and lines represent predicted inhibition in the network. (C) IPA analysis of diseases and biological functions with the MD unique genes. Orange colored bars represent signaling pathways that are not common with JPS as shown in figure 4A. (D) Validation of target gene expression of orange colored signaling pathways in figure 5C with quantitative real-time PCR (qRT-PCR). *CDX2*, a downstream effector of CDX gastrointestinal signaling (left) and *HES1*, a downstream effector of Notch signaling (middle) show trends of increased expression in MD, however, it is not statistically significant. Expression of *GLI1*, a downstream effector of Sonic hedgehog signaling (right), is significantly increased in MD (* means P value<0.05).

### Role of Hedgehog signaling in the pathogenesis of MD

It has been previously reported that Hh signaling plays essential roles in stomach development and normal homeostasis as well as carcinogenesis in the stomach (Jian-Hui, Er-Tao et al. 2016, Fattahi, Nikbakhsh et al. 2021, Akhtar, Maqbool et al. 2022). We used the MD mouse model metallothionein (MT)-TGF-α to examine the role of Hh signaling in the pathogenesis of MD (Dempsey, Goldenring et al. 1992, Goldenring, Poulsom et al. 1996, Nomura, Settle et al. 2005). MT-TGF-α mice were treated with either a Hh signaling inhibitor, sonidegib (5 mg/kg) or DMSO control via intraperitoneal (i.p.) injection every other day for three weeks (Figures 6A). Sonidegib treatment did not affect weight or wellbeing of the mice. We observed that the Hh downstream effector molecule GLI1 showed decreased expression after sonidegib treatment by immunoblot, confirming the effect of Hh signaling inhibition by sonidegib (Figure 6B). Hh inhibition partially rescued the MD phenotypes in MT-TGF-α mice. Numbers of parietal and chief cells were increased (Figure 6C and 6E), number of foveolar cells were decreased, and number of neck cells were not significantly changed (Figure 6D and 6F). These findings suggest that Hh signaling plays an important role in the pathogenesis of MD and that it may be a potential therapeutic target.

**Figure 6.**
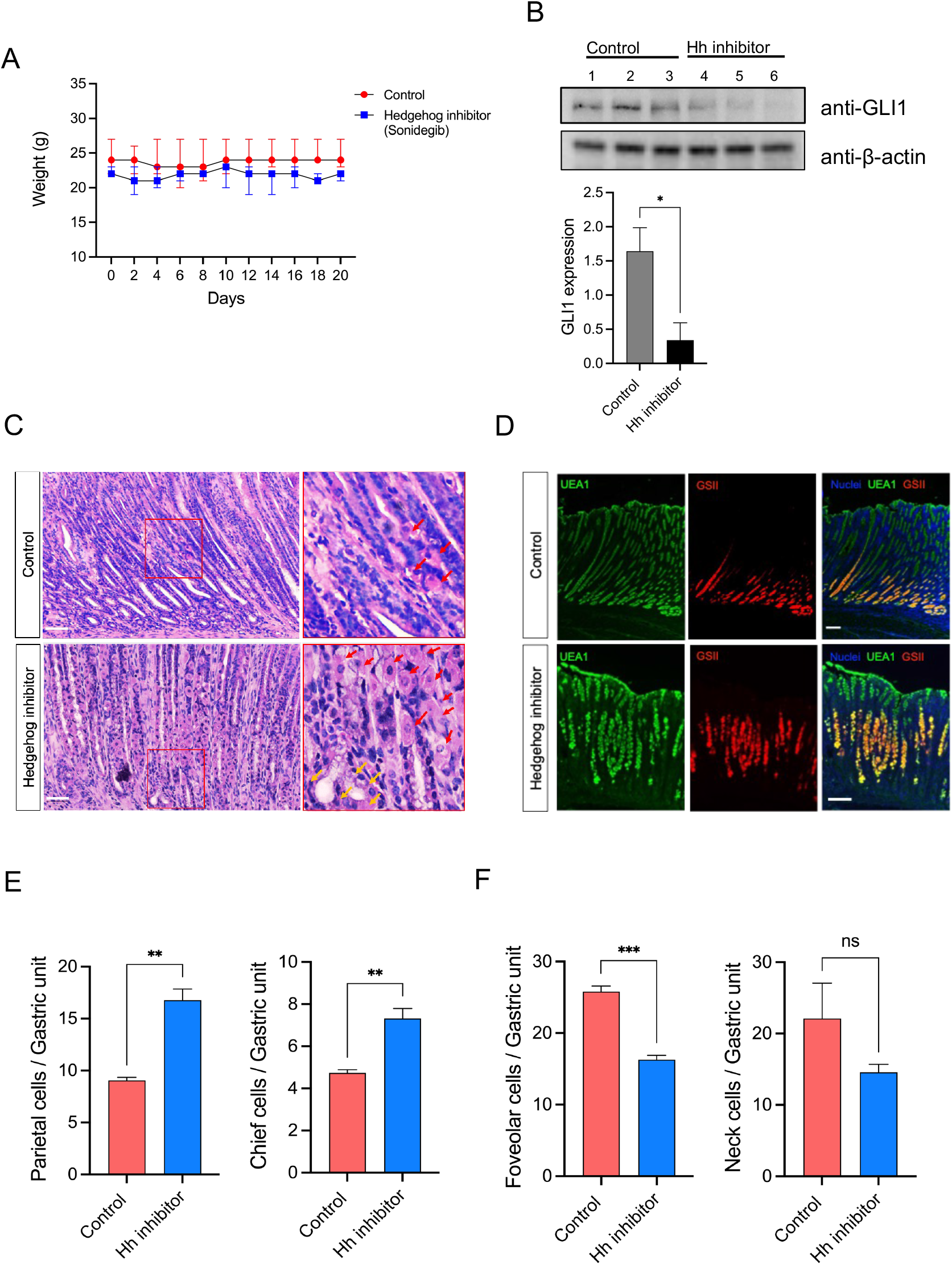
Role of hedgehog signaling in the pathogenesis of MD. Transgenic mice overexpressing TGF-α (MD mouse model MT-TGFA) were treated with a hedgehog (Hh) inhibitor (sonidegib) or control (DMSO) treatment with 5 mg/kg dose on intraperitoneal (i.p.) injection for three weeks. (A) The body weights of each experimental group (sonidegib, n=3 and control, n=3) were measured every other day for 21 days. Sonidegib or DMSO treatment did not affect the weight of mice. (B) Immunoblots with anti-GLI1 and anti-beta-Actin antibodies show that Hh signaling is effectively inhibited by the sonidegib treatment. Bar graph represents quantification of the immunoblots (* means P value<0.05). (C and E) Hematoxylin and eosin (H&E) stains reveal that sonidegib treatment increases the number of parietal cells (red arrows) and chief cells (yellow arrows) in MT-TGFA mice. Scale bars represent 50 µm. The positive cells from each gastric unit were detected by QuPath software (ver.0.4.3) under their algorithm and calculated the average number of cells with SEM (standard error of the means) were visualized. The differential numbers of each cell type were analyzed with unpaired t-test by GraphPad (Prism10) software. Numbers of parietal and chief cells were counted by the QuPath software (** means P-value <0.01). (D and F) UEA1 positive foveolar cells are significantly decreased by sonidegib treatment. Number of GSII positive neck cells shows decreasing trend, but it is not statistically significant. UEA1 and GSII positive cells were counted using the QuPath software. Scale bars represent 100 µm. *** means P-value <0.001 and ns not significant.

## DISCUSSION

The histomorphological characteristics of transforming growth factor alpha (TGF-α) overexpression include oxyntic atrophy (loss of parietal cells) and massive foveolar hyperplasia in the gastric body/fundus of patients with MD as well as transgenic mice overexpressing TGF-α (Goldenring, Poulsom et al. 1996, Nomura, Settle et al. 2005). MD is a rare disease and can be challenging to diagnose since there are many mimickers that share similar histopathologic features with MD (Rich, Toro et al. 2010, Huh, Coffey et al. 2016). In our prior clinical trial assessing the effects of cetuximab for the treatment of MD, juvenile polyposis syndrome (JPS) was one of the most common diseases that was misdiagnosed as MD. In the study, we analyzed the RNA sequencing (RNA-seq) datasets of MD, JPS, and normal (NL) samples and identified that claudin 18 (CLDN18) is a diagnostic marker for MD.

CLDN18, a member of the claudin protein family, is one of the structural components of tight junctions. It has two splice variants, claudin 18.1 (CLDN18.1) and claudin 18.2 (CLDN18.2), which are specifically expressed in lung and stomach, respectively. Recently, CLDN18 has emerged as a therapeutic target in advanced gastric cancer. In gastric cells, claudin 18.2 expression was regulated through the protein kinase C (PKC)/ mitogen-activated protein kinase (MAPK)/ activator protein (AP)-1 dependent pathway (Yano, Imaeda et al. 2008). Supportively, the claudin 18 positive cell lines associated with the proliferative properties, indicating the increased Ki-67 labeling at the invasive gastric cancer (Oshima, Shan et al. 2013). Recent metanalysis on the CLDN18.2 expression in gastric cancer did not show significant correlation between CLDN18.2 expression and pathological or prognosis features such as TNM stages, HER2, grading or overall survival. However, the authors suggested the relationship between CLDN18.2 and pathological/prognosis features will depend on the percentage of staining of tumor cells (Ungureanu, Lungulescu et al. 2021). Indeed, clinical trials of zolbetuximab, a monoclonal antibody against CLDN18.2, showed the level of CLDN18.2 expression was associated with the response of gastric cancer to the treatment (Singh, Toom et al. 2017, Dottermusch, Kruger et al. 2019, Athauda and Chau 2021). In the current study, the comparative analysis of RNA-seq showed that *CLDN18* is upregulated in MD when compared to JPS. We showed that CLDN18.2 expression was higher in MD stomachs than in the JPS group by IHC. Even though this finding needs to be confirmed with more MD and JPS tissue samples, it suggests that CLDN18.2 can be used as a diagnostic marker that can distinguish MD from JPS. Since zoltuximab is an effective gastric cancer treatment in tumors with high CLDN18.2 expression, it also needs to be investigated if this therapy would be similarly effective in MD. Both MD and JPS are premalignant diseases that show similar histopathologic findings. We presumed that commonly expressed genes would account for the similarity, and we analyzed these mutual genes using IPA^®^ canonical pathway analysis. It revealed that MD and JPS shared many signaling pathways including those associated with ERBB activation such as ERK/MAPK signaling, PI3K/AKT signaling, and PKA signaling, as well as Pre-Notch process signaling and mTOR signaling pathways. ERBB signaling is activated in MD because ERBB1/EGFR ligand TGF-α is overexpressed. Although JPS is caused by loss-of-function mutations in genes involved SMAD4 signaling, our data suggests there might be crosstalk between SMAD4 and ERBB signaling pathways in JPS which results in the activation of ERBB signaling which could cause similar histopathologic features in MD and JPS. Activation of ERBB, Notch, and mTOR signaling pathways have been associated with gastric cancer (Ma, Li et al. 2022, Song, Gao et al. 2023). This suggests that activation of ERBB, Notch, and mTOR signaling pathways might similarly lead to cancer progression in MD and JPS.

MD is a very rare disease, and little is known about the pathogenesis except for the role of EGFR activation through TGF-α overexpression. Comparative analysis between MD and NL revealed that sonic hedgehog (SHH) signaling is highly associated with MD, and SHH expression and activation of the SHH pathway have been associated with the prognosis of gastric cancer (Kim, Ko et al. 2012, Saze, Terashima et al. 2012). In the current study, we found that treatment with an FDA-approved Hedgehog (Hh) inhibitor, sonidegib, could partially rescue the pathological phenotypes in a MD mouse model. This suggests the Hh signaling pathway plays an important role in the pathogenesis of MD and it may be a novel therapeutic target in treating the disease. Due to the rarity of MD diagnosis, the pathogenesis of the disease is poorly understood. Many disease entities including JPS can mimic histopathologic features of MD, making an accurate diagnosis challenging. Here, we performed RNA-seq of MD, JPS, and NL stomach samples, and identified that CLDN18.2 is a diagnostic biomarker that can help distinguish MD from JPS, its most common mimicker. We also found that Hh signaling is essential in the pathogenesis of MD and it could be used as a novel therapeutic target. Further studies are needed to confirm the findings in the current study.

## Funding

This work was supported, in part, by National Institute of Diabetes and Digestive and Kidney (NIDDK) grant K08DK124686 (to W.J.H).

## Notes

### Competing Interest Statement

The authors have declared no competing interest.

